# The SET1 protein of *Leishmania donovani* moderates the parasite’s response to a hostile oxidative environment

**DOI:** 10.1101/2023.09.21.558927

**Authors:** Jyoti Pal, Varshni Sharma, Arushi Khanna, Swati Saha

**Affiliations:** Department of Microbiology University of Delhi South Campus New Delhi-110021 INDIA

**Keywords:** *Leishmania donovani*, trypanosomatids, SET domain, SET proteins, oxidative stress, protozoan parasite

## Abstract

SET domain proteins mediate their effects through the methylation of specific lysine residues on target substrates, resulting in either the stimulation or repression of downstream processes. Initially identified as histone lysine methyltransferases, they are now known to target a wide-ranging conglomeration of non-histone subtrates as well. Twenty-nine SET domain proteins have been identified in *Leishmania donovani* through genome sequence annotations. This study initiates the first investigation into these proteins. We find LdSET1 is predominantly cytosolic and constitutively expressed. The *set1* gene is not essential, although its deletion slows promastigote growth and hypersensitizes the parasite to hydroxyurea-induced G1/S arrest. Intriguingly, *set1*-nulls proliferate more proficiently than *set1^+/+^* parasites within host macrophages, suggesting that LdSET1 moderates the parasite’s response to the inhospitable intracellular oxidative environment. *set1*-null parasites are highly tolerant to H_2_O_2_-induced oxidative stress in *in vitro* promastigote cultures as well, reflected in their growth pattern as well as almost complete absence of DNA damage at the H_2_O_2_ concentrations tested. This is linked to ROS levels remaining virtually unperturbed in response to H_2_O_2_ treatment in *set1*-nulls, contrasting to increased ROS in *set1^+/+^* cells under similar conditions. In analyzing the cell’s ability to scavenge hydroperoxides we find that peroxidase activity is not upregulated in response to H_2_O_2_ exposure in *set1*-nulls. Rather, the constitutive basal levels of peroxidase activity are significantly higher in these cells, implicating this to be a factor contributing to the parasite’s apparent resistance to H_2_O_2_. The higher levels of peroxidase activity in *set1*-nulls are coupled to upregulation of tryparedoxin peroxidase transcripts in these cells. Thus, LdSET1 tunes the parasite’s behavior within host cells, enabling establishment and persistence of infection, and maintaining the balance with host without eradicating the host cell population it needs for survival. These findings uncover a new dimension to the mechanisms controlling the interplay of *Leishmania*-host interactions.

**Author Summary:** Leishmaniases are a group of diseases afflicting people across 88 countries. Manifested in three forms: cutaneous, subcutaneous and visceral, different species cause the different forms of the disease. *Leishmania donovani* is one of the species causing Visceral Leishmaniasis, and its cellular processes are an area of intensive investigations. These protozoan parasites display an unusual pattern of transcription, and gene regulation occurs primarily through epigenetic mechanisms and post-transcriptional processes. SET proteins control gene expression by methylating histones as well as non-histone proteins. In this study we have examined the role of the LdSET1 protein. We find that while the *set1* gene is not essential to cell survival, deletion of the *set1* gene makes the parasite highly tolerant to the effects of an oxidizing environment, signifying that the LdSET1 protein plays a role in regulating the cell’s response to oxidative stress. LdSET1 appears to moderate the parasite’s survival in the oxidative intracellular milieu of host macrophages, controlling it such that infection is firmly established, and persistent, without wiping out the population of host cells it needs for survival.

## Introduction

The SET domain proteins get their name from the three Drosophila proteins the domain was first identified in: **<u>s</u>**uppressor of variegation [**<u>S</u>**u(var)3-9], **<u>e</u>**nhancer of zeste [E(z)], **<u>t</u>**rithorax (reviewed in [1]). Initially identified as proteins that methylate histones at specific lysine residues, SET proteins were subsequently found to target non-histone substrates as well, and can mediate mono-, di- and/or tri-methylation of their target residues. Common histone targets include H3K4, H3K9, H3K27, H3K36 and H4K20 (reviewed in [1–3]). The impacts of these histone methylation events vary widely, and thus while H3K4 methylation is associated with transcriptional upregulation, H3K9 and H3K27 methylation marks mediate transcriptional repression. Non-histone substrates include tumor suppressors (such as p53 and Rb), transcription factors (like GATA4 and TAF_10_) and signalling proteins (such as STAT3) (reviewed in [1–3]). SET domain proteins modulate a wide range of cellular processes, such as DNA replication, DNA repair, transcriptional activation as well as silencing, mRNA splicing, heterochromatin formation, and X-chromosome inactivation. Abrogation of SET domain protein functions have been linked to various types of cancer [1, 3].

While exhaustively studied in yeast and mammalian cells, limited information is available about protozoan SET proteins. The most widely investigated are the *Plasmodium falciparum* SET proteins, where six of the ten proteins identified appear to be essential for survival of blood stage parasites. The histone target residues of six PfSET proteins have been identified, and their functional roles characterized to varying degrees [4–6]. The *Tetrahymena* EZL3 SET protein has been found to modulate development and progeny viability [7]. The non-nuclear *Toxoplasma* SET domain protein modulates host cell invasion as well as the parasite’s exit from the host cell [8]. *In vitro* biochemical assays have uncovered the target residues of three SET domain proteins identified in *Entamoeba histolytica*, two of which appear to be involved in phagocytosis [9].

*Leishmania* species are trypanosomatids that are endemic to 88 countries, and according to WHO around 12 million people are at risk globally (https://www.who.int/news-room/fact-sheets/detail/leishmaniasis). Leishmaniases are manifested in three forms: cutaneous, subcutaneous and visceral, and different *Leishmania* species cause the different forms of the disease. In South Asia the prevalent form of the disease is visceral leishmaniasis (VL) or kala-azar, caused by *Leishmania donovani.* VL is treatable, but treatment regimens are lengthy, expensive, and have toxic side effects. Adding to that the problems of growing drug resistance, risks due to HIV-*Leishmania* co-infection, and emerging threat of PKDL (post kala-azar dermal leishmaniasis) in the Indian subcontinent, this parasite’s cellular biology remains an area of interest to several research groups around the world. While no work regarding SET domain proteins in *Leishmania* species has been reported, twenty-nine SET domain proteins have been identified in *Trypanosoma brucei* by whole genome sequence annotations, and orthologs of all twenty-nine are identifiable in *L. donovani*. The subcellular localization of the *T. brucei* SET proteins has been examined in blood form (BF) as well as procyclic form (PF), by epitope-tagging coupled to immunofluorescence [10–12]. Eight of the twenty-nine proteins localize to the nucleus in the BF parasites, though not exclusively so. Four of these eight proteins localize to the nucleus in the PF parasites as well. Six SET proteins localize to the nucleus (though not exclusively) in the PF parasites only. Two proteins, TbSET26 and TbSET27, have been found to be enriched at the Transcriptional Start Regions (TSRs) [12]. Furthermore, TbSET27 has been found to exist as part of a multi-protein complex (SPARC) that modulates the accuracy of transcription initiation at the TSRs [13].

This report presents the first data regarding SET proteins across *Leishmania* species, and directly investigates the functional role of SET1. Our study reveals that the *set1* gene of *L. donovani* is not essential for cell survival. The LdSET1 protein is found to play a role in mediating the parasite’s response to an oxidative environment in both, promastigotes (the extracellular form in insect host) and amastigotes (the intracellular form in mammalian host).

## Results

### Leishmania donovani SET1 is constitutively expressed and is predominantly cytosolic

In the absence of genome sequence information of *L. donovani* 1S (Ld1S), the *set1* gene (∼1.42 kb) of Ld1S was cloned by amplification off genomic DNA using primers designed against its ortholog in *L. donovani* BPK282A1 (LdBPK_360230.1), whose sequence was obtained from TriTrypDB ([14] www.tritrypdb.org). The cloned amplicon was sequenced to confirm its veracity (GenBank Accession no: OR479702). It was found to carry only one SNP in comparison with the LdBPK_360230.1 gene, which resulted in a change from leucine to valine. A comparison of the derived amino acid sequence of LdSET1 with those of other trypanosomatid SET1 proteins using Clustal Omega analysis [15] and blastp (https://blast.ncbi.nlm.nih.gov.in) revealed that LdSET1 showed 35-40% identity over a coverage of 91% with SET1 of *Trypanosoma* species, and 90-99% identity with SET1 of other *Leishmania* species over 100% coverage (S1 Fig). Analysis of the SET1 amino acid sequence revealed that the ∼51.5 kDa protein carries a SET domain between amino acids 304-422, and a post-SET domain at the C-terminal end between residues 430-446 (S2A Fig). SET domains are often flanked by pre-SET and post-SET domains. While the pre-SET domain is believed to overall help stabilize the SET protein structure via its interactions with the residues of the SET domain, the post-SET domain forms part of the active site hydrophobic channel [16].

The *set1* gene was expressed episomally in fusion with a C-terminal FLAG tag in *L. donovani* promastigotes (the extracellular flagellate insect form; S2B Fig), and the subcellular localization of LdSET1-FLAG analyzed by indirect immunofluorescence as described in Methods, using kinetoplast morphology and segregation pattern as cell cycle stage marker [17]. LdSET1-FLAG was found to be primarily cytosolic at all stages of the cell cycle (S2C Fig). The expression of LdSET1 in logarithmically growing as well as stationary phase *L. donovani* promastigotes was analyzed in whole cell extracts using anti-LdSET1 antibodies already available in the lab, and found to be comparable at both stages (S3A Fig). Examination of whole cell lysates isolated from procyclics (early stage promastigotes normally existing in insect midgut) and metacyclics (late stage promastigotes found usually in insect salivary glands; the infective stage) revealed that expression was more or less equivalent in both types of promastigotes (S3B Fig). Upon examining LdSET1 expression at different stages of the cell cycle using promastigotes that had been synchronized at the G1/S boundary with a hydroxyurea block and then released into S phase (described in Methods), it was observed that LdSET1 expression was maximum in S phase (S3C Fig). This was in contrast to expression of another *L. donovani* SET protein, LdSET29, which was expressed more or less equivalently at all stages of cell cycle (S3C Fig). Collectively, these data demonstrate that LdSET1 is robustly expressed in promastigotes (in both procyclics and metacyclics), and is constitutively expressed, predominantly in the cytoplasm, with maximal expression detectable in S phase.

### LdSET1-depleted parasites display greater competency in survival within host macrophages than wild type parasites

To investigate the role of SET1 in the *Leishmania* cell we replaced the *set1* genomic alleles sequentially by homologous recombination (as detailed in Methods). The first allele was replaced with a hygromycin resistance cassette, and the authenticity of recombination at both ends verified by PCRs across the deletion junctions (S4 Fig). Western blot analysis revealed a corresponding decrease in the levels of the LdSET1 protein (Fig 1A). Analysis of the growth pattern of these *set1-*heterozygous knockout (*set1*^-/+^) promastigotes revealed that the observed reduction in LdSET1 expression upon knockout of one *set1* allele did not have any impact on growth and survival under normal conditions (Fig 1B). The cell cycle progression pattern of *set1*^-/+^ cells, synchronized at the G1/S boundary with hydroxyurea (HU) and then released into S phase (as described in Methods), was similar to that of *set1*^+/+^ cells (Fig 1C). To examine the effect of loss of one *set1* allele on amastigote growth and propagation, we infected *set1*^-/+^ metacyclics into murine macrophage cells J774A.1 (as described in Methods) and followed their survival over 48 hours after infection. As seen in Fig 1D, the intracellular load of *set1*^-/+^ parasites was comparable to that of *set1*^+/+^ parasites at the end of the 5 hour parasite-host cell incubation period (5 hour time-point), signifying that the ability to infect host cells *per se* was not altered in *set1*^-/+^ cells. However, unexpectedly, *set1*^-/+^ parasites survived and proliferated more competently than *set1*^+/+^ parasites within the host cells, as determined by the higher intracellular parasite load 24 hours and 48 hours after infection (Fig 1D). Thus, the levels of SET1 protein in *set1^-/+^* parasites are sufficient for the promastigotes to remain unperturbed under normal *in vitro* growth conditions, but insufficient to maintain the normal parasite-host cell balance in case of the amastigotes.

**Fig 1.**
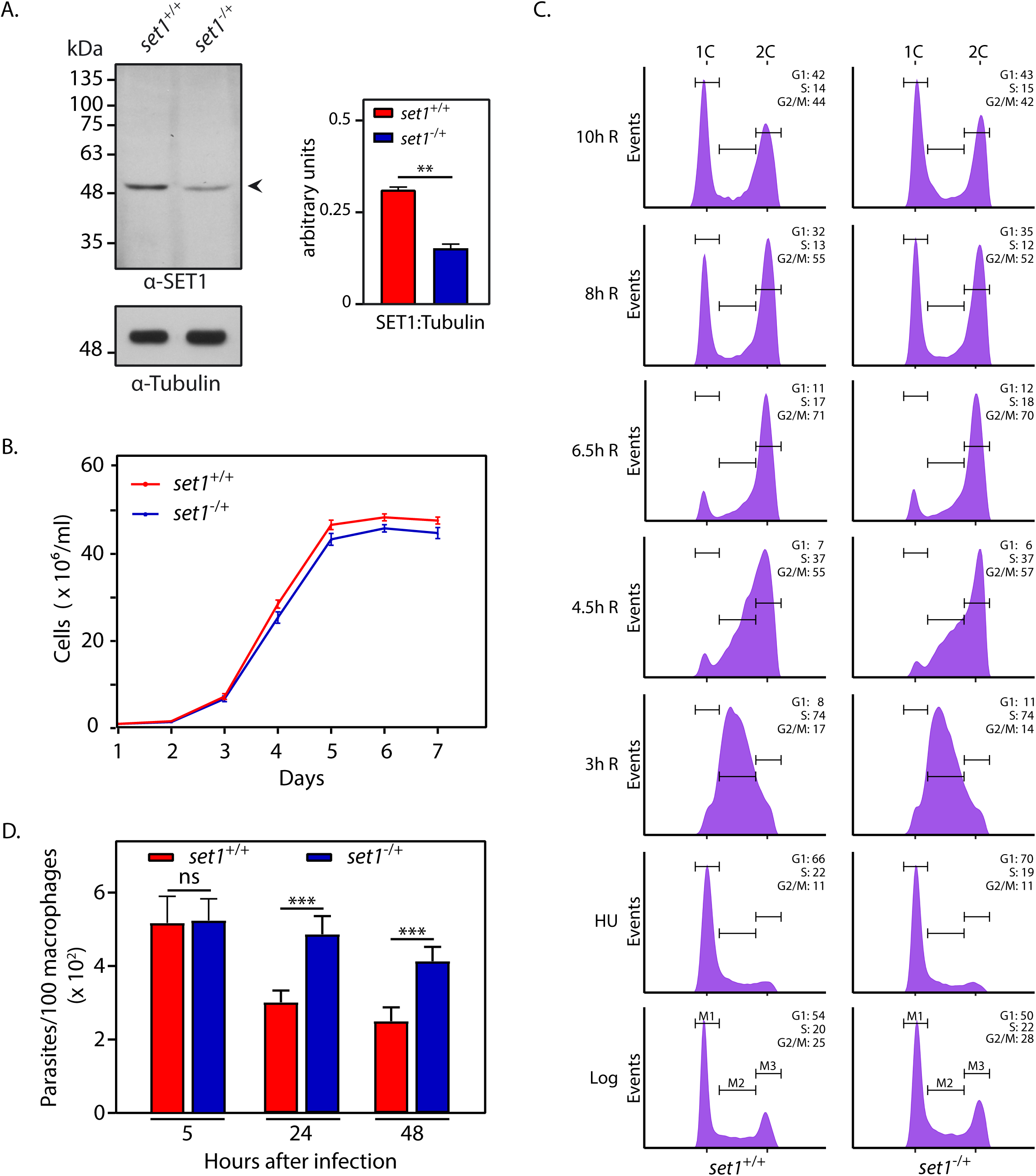
Effect of LdSET1 depletion on parasite growth and cell cycle progression pattern. A. Western blot analysis of whole cell extracts isolated from 1×10^8^ logarithmically growing promastigotes using anti-SET1 antibodies (1:2500 dilution). Loading control: tubulin. Quantitation was carried out using ImageJ analysis and plotted values represent average of three experiments. Error bars represent standard deviation and statistical significance was determined using student’s *t*-test. ** represents *p* value <0.005. **B.** Analysis of growth of parasites. Values plotted represent average of three experiments. Each experiment was performed with two technical replicates. Error bars represent standard deviation. **C.** Flow cytometry analysis of HU-synchronized promastigotes. Time-points at which cells were sampled are indicated on the left of each row of histograms. R: release (3h R signifies 3 hours after release from HU-induced block). 30000 events were analyzed at every time-point. M1, M2 and M3 represent gating for cells in G1, S and G2/M respectively. Percent cells in each cell cycle phase are indicated in upper right-hand corner of each histogram. The experiment was done thrice, with comparable results, and data of one experiment is shown. **D.** Analysis of parasite survival in host macrophages. Z-stack imaging using confocal microscopy was employed to count the number of intracellular parasites (DAPI-stained nuclei served as marker). The experiment was performed thrice and average values are plotted in the bar graph. Error bars depict standard deviation and student’s *t*-test was used to assess statistical significance. ***: *p* value <0.0005. ns: statistically not significant. *set1^+/+^* cells: Ld1S::hyg cells. *set1^-/^*^+^ cells: *set1-*heterozygous knockout cells.

### LdSET1-depleted promastigotes are more tolerant to an oxidative environment than wild type promastigotes

Considering the oxidative intracellular environment of macrophage host cells we examined the growth of LdSET1-depleted promastigotes in conditions where oxidative stress has been induced. Thus, *set1*^-/+^ parasite cultures were initiated from stationary phase cultures, and hydrogen peroxide varying over 25-200 µM was added to the cultures on Day 3 (as described earlier). The parasites were incubated in this H_2_O_2_-containing medium (M199) for 5 hours before replacing it with fresh H_2_O_2_-free medium and continuing incubation. Growth was monitored over the next few days, and on comparing the behaviour of *set1*^-/+^ promastigotes with that of *set1*^+/+^ promastigotes it was observed that while neither cell type displayed any obvious response to lower concentrations of H_2_O_2_ (25-50 µM), they behaved quite differently in response to higher H_2_O_2_ concentrations. *set1*^+/+^ promastigotes grew slower in response to 100 µM H_2_O_2_ treatment and displayed severely compromised growth in response to 200 µM H_2_O_2_ treatment; in contrast, *set1*^-/+^ parasites displayed a very moderate response to 100 µM H_2_O_2_ treatment, and were able to recover and continue to grow well in response to 200 µM H_2_O_2_ treatment as well (Fig 2A). The altered behaviour of LdSET1-depleted parasites upon exposure to H_2_O_2_ was more apparent when parasites were incubated in 100 µM H_2_O_2_ over several days (with replenishment of H_2_O_2_ every 10 hours), wherein growth of *set1*^+/+^ cells was distinctly compromised while *set1*^-/+^ cells continued to thrive (Fig. 2B). These data indicate that SET1 regulates the parasite’s response to an oxidative milieu under *in vitro* conditions as well.

**Fig 2.**
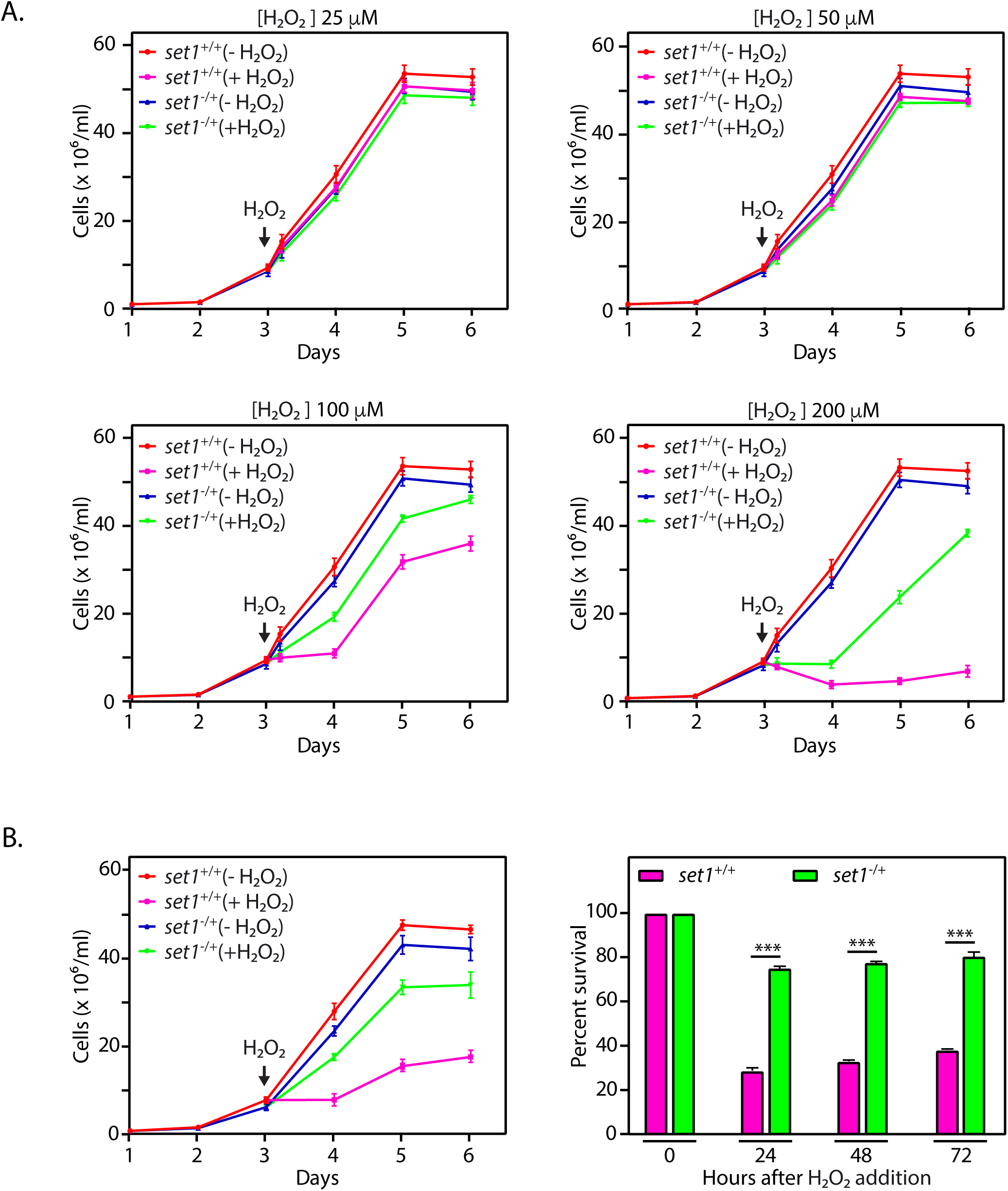
Effect of LdSET1 depletion on promastigote growth and survival in response to H_2_O_2_ exposure. **A.** Cultures were initiated from stationary phase cultures, and 0-200 μM H_2_O_2_ was added to the cultures 48 hours after initiation. At the time of addition of H_2_O_2_ the cultures were split into two, with half the cells receiving H_2_O_2_ treatment and the other half being carried forward as the untreated culture. After a 5-hour exposure to H_2_O_2,_ the parasites were re-fed with fresh H_2_O_2_-free medium, and cells counted every 24 hours. Graphs show growth patterns of the cells. The experiment was done thrice, with two technical replicates in each experiment. Values plotted are average of three experiments and error bars depict standard deviation. The experimental data has been split into four panels for easier viewing and thus the (-H_2_O_2_) data is identical in all four panels. **B.** Cultures were initiated from stationary phase cultures, and 100 μM H_2_O_2_ was added to the cultures 48 hours after initiation. At the time of addition of H_2_O_2_ the cultures were split into two, with half the cells receiving H_2_O_2_ treatment and the other half being carried forward as the untreated culture. H_2_O_2_-treated cultures were maintained in H_2_O_2_-containing medium, with H_2_O_2_ being replenished every 10 hours, and cells were counted every 24 hours. Left panel: Growth analysis of the cultures. Right panel: Survival percent of H_2_O_2_-treated cultures with reference to untreated cultures, was determined by dividing the number of cells in treated cultures by the number of cells in untreated cultures and multiplying by 100. The experiment was done thrice, with two technical replicates in each experiment. Values plotted are average of three experiments and error bars depict standard deviation. Student’s *t*-test was used to assess statistical significance. ***: *p* value <0.0005. *set1^+/+^* cells: Ld1S::hyg cells. *set1^-/^*^+^ cells: *set1-*heterozygous knockout cells.

### The set1 gene is not essential for cell survival

The second *set1* allele was replaced with a *neo^r^* cassette, and genuine recombination at the 3’end checked by PCRs across the deletion junction while the authenticity of recombination at the 5’end was checked by inverse PCRs across the deletion junction (S5 Fig). Western blot analysis confirmed the absence of LdSET1 expression in *set1*-nulls (*set1*^-/-^, Fig 3A), and analysis of growth revealed that promastigotes grew slower when completely devoid of LdSET1, entering log phase three days later than *set1*^+/+^ cells and never reaching the same cell density as *set1*^+/+^ cells before entering stationary phase (Fig 3B). The generation time of *set1*^-/-^ cells was found to be considerably longer than that of *set1*^+/+^ cells (∼23 hours as compared to the usual ∼9.7 hours; Fig. 3C). Examination of cell cycle progression patterns by flow cytometry analysis of HU-synchronized promastigotes revealed that *set1*^-/-^ cells displayed a heightened sensitivity to HU-induced G1/S arrest, with a large fraction of the cells failing to be released into S phase upon removal of HU (Fig. 3D). While the fraction of cells that got released from the HU-induced block appeared to traverse S phase and G2/M in a manner comparable to control cells, no definitive conclusion regarding the cause for increased generation time could be drawn from this data. It is possible that the span of G1 phase is longer in *set1^-/-^* cells; this needs further investigation. From the data in Figs. 1B-C and 3B-D, we concluded that under normal *in vitro* growth conditions while a partial depletion of LdSET1 (to ∼50% wild type levels) did not have any impact on growth and cell cycle progression, complete elimination of LdSET1 expression slowed down growth and increased the generation time more than two-fold. The survival of the parasite in the absence of the *set1* gene indicates that *set1* is not essential to the parasite.

**Fig 3.**
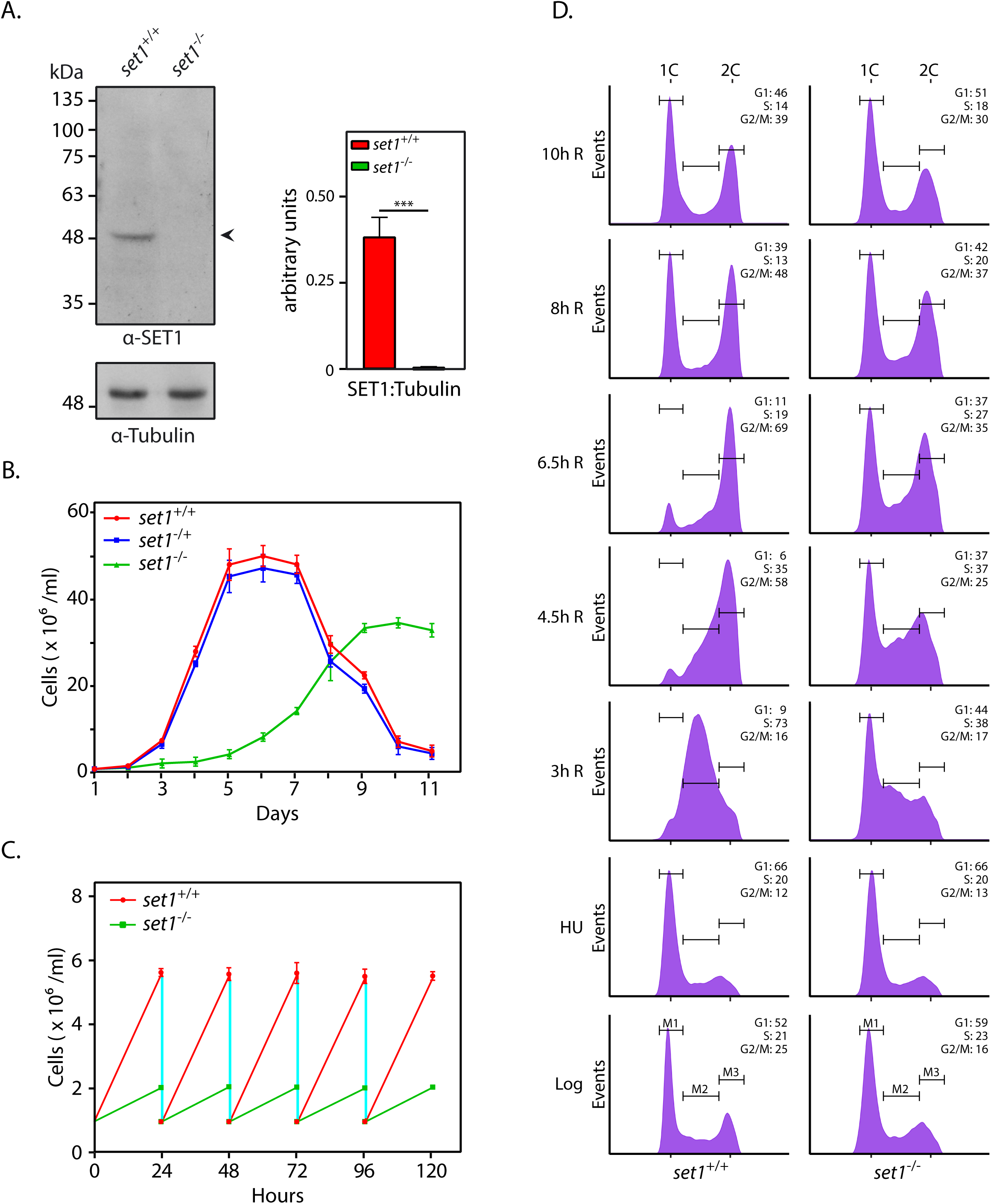
Effect of elimination of *set1* gene on promastigote growth and cell cycle progression. **A.** Western blot analysis of whole cell extracts isolated from 1×10^8^ logarithmically growing promastigotes using anti-SET1 antibodies (1:2500 dilution). Loading control: tubulin. Quantitation was carried out using ImageJ analysis and plotted values represent average of three experiments. Error bars represent standard deviation and statistical significance was determined using student’s *t*-test. *** represents *p* value <0.0005. **B.** Analysis of growth of parasites. Cultures were initiated at 1×10^6^ cells/ml, from stationary phase cultures. Values plotted represent average of three experiments. Each experiment was performed with two technical replicates. Error bars represent standard deviation. **C.** Generation time of cells was determined by initiating cultures from logarithmically growing cultures, at 1×10^6^ cells/ml, and diluting the cultures to 1×10^6^ cells/ml every 24 hours after counting them. The experiment was done thrice and average values are plotted, with error bars depicting standard deviation. **D.** Flow cytometry analysis of HU-synchronized promastigotes. Time-points at which cells were sampled are indicated on the left of each row of histograms. 30000 events were analyzed at every time-point. M1, M2 and M3 represent gating for cells in G1, S and G2/M respectively. Percent cells in each cell cycle phase are indicated in upper right-hand corner of each histogram. The experiment was done thrice, with comparable data, and data of one experiment is shown. *set1^+/+^* cells: Ld1S::neo-hyg cells. *set1^-/^*^+^ cells: *set1-*heterozygous knockout cells. *set1^-/^*^-^ cells: *set1*-nulls.

### Set1-nulls survive and propagate more proficiently than wild type cells within host macrophages

Even more interestingly, these slower growing *set1*^-/-^ parasites (Fig 3B-C) survived and propagated more proficiently than *set1*^+/+^ parasites within host macrophages, though not to the same extent as *set1*^-/+^ parasites, reflecting the longer generation time of *set1*^-/-^ cells as compared to *set1*^-/+^ cells (Fig 4A). Thus, LdSET1 appears to be moderating the parasite’s response to the inhospitable intracellular environment of host cells. The differential response of *set1*^-/-^ promastigotes to an oxidative growth environment *in vitro* was even more pronounced. While *set1*^-/-^ cells in general grew slower than *set1*^+/+^ promastigotes, the parasites did not demonstrate any apparent response to a five-hour H_2_O_2_ treatment over concentrations of 25-200 µM (Fig 4B), and nor was any response evident when the parasites were incubated in 100 µM H_2_O_2_ over several days (Fig 4C). When treated with H_2_O_2_ over concentrations of 500-1000 µM for five hours, however, *set1*-nulls did not survive (S6A Fig). Elimination of LdSET1 from the cell thus appears to make the parasites extremely tolerant to the effects of an H_2_O_2_-induced oxidative environment.

**Fig 4.**
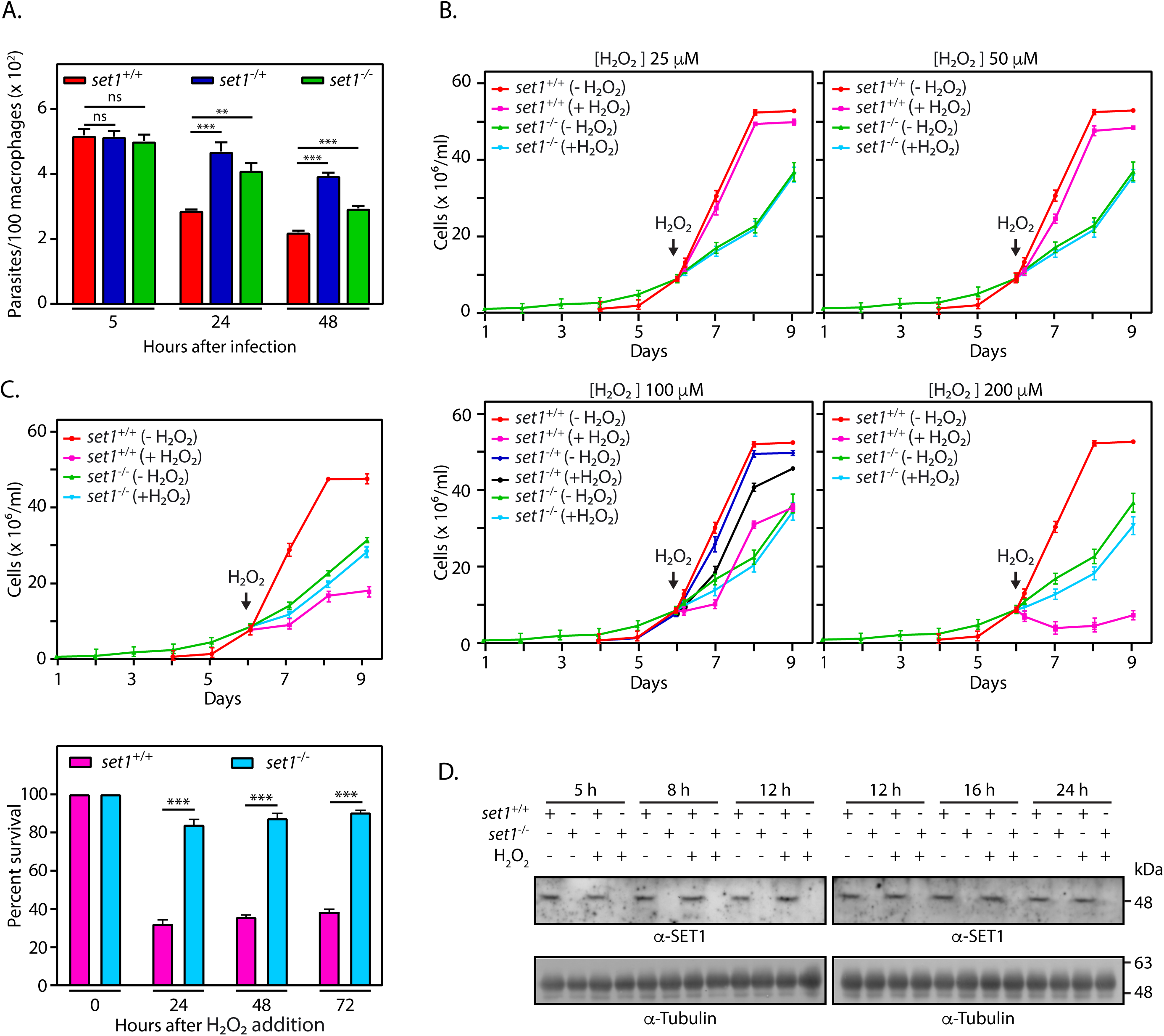
Effect of oxidizing environment on *set1*-null parasites. **A.** Analysis of parasite survival in host macrophages. Z-stack imaging using confocal microscopy was employed to count the number of intracellular parasites (DAPI-stained nuclei served as marker). The experiment was performed thrice and average values are plotted in the bar graph. Error bars depict standard deviation and student’s *t*-test was used to assess statistical significance. ***: *p* value <0.0005, **: *p* value <0.005, ns: statistically not significant. **B.** Effect of H_2_O_2_ exposure on promastigote growth and survival. Cultures were initiated from stationary phase cultures, and 0-200 μM H_2_O_2_ was added to the cultures at 7-9 x10^6^ cells/ml. At the time of addition of H_2_O_2_ the cultures were split into two, with half the cells receiving H_2_O_2_ treatment and the other half being carried forward as the untreated culture. Parasites were re-fed with fresh H_2_O_2_-free medium after a 5 hour-exposure to H_2_O_2_, and cells counted every 24 hours. The experiment was done thrice, with two technical replicates in each experiment. Values plotted are average of three experiments and error bars depict standard deviation. The experimental data has been split into four panels for easier viewing and thus the (-H_2_O_2_) data is identical in all four panels. **C.** Effect of prolonged H_2_O_2_ exposure on promastigote growth and survival. Cultures were initiated from stationary phase cultures, 100 μM H_2_O_2_ added to the cultures at cell density 7-9 x10^6^ cells/ml (at the time of addition of H_2_O_2_ the cultures were split into two, with half the cells receiving H_2_O_2_ treatment and the other half being carried forward as the untreated culture), and cultures maintained in H_2_O_2_-containing medium (with H_2_O_2_ being replenished every 10 hours), with cells being counted every 24 hours. Upper panel: Growth analysis of the cultures. Lower panel: Survival percent of H_2_O_2_-treated cultures with reference to untreated cultures, determined by dividing the number of cells in treated cultures by the number of cells in untreated cultures and multiplying by 100. The experiment was done thrice, with two technical replicates in each experiment. Values plotted are average of three experiments and error bars depict standard deviation. Student’s *t*-test was used to assess statistical significance. ***: *p* value <0.0005. **D.** Western blot analysis of whole cell extracts isolated from 8×10^7^ cells that were exposed to 100 μM H_2_O_2_ for 5 hours, using anti-SET1 antibodies (1:1000 dilution). Loading control: tubulin. Time-points indicate hours after start of the H_2_O_2_ exposure. *set1^+/+^* cells: Ld1S::neo-hyg cells. *set1^-^ _/_*_+_ cells: *set1-*heterozygous knockout cells. *set1^-/^*^-^ cells: *set1*-nulls.

The effect of H_2_O_2_ treatment on LdSET1 expression in *L. donovani* promastigotes was analyzed by incubating cells with H_2_O_2_ (100 µM) for 5 hours, followed by continued growth in H_2_O_2-_free medium, and isolation of whole cell lysates at various time intervals thereafter. The lysates were analyzed using western blotting with anti-SET1 antibodies, and as seen in Fig 4D, LdSET1 expression levels did not change in response to H_2_O_2_ treatment. To see if H_2_O_2_ treatment affected the subcellular localization of LdSET1, *L. donovani* promastigotes expressing LdSET1-FLAG were treated with H_2_O_2_ (100 µM) for 5 hours, collected by centrifugation, and analyzed by indirect immunofluorescence. No change in subcellular localization was apparent (S6B Fig). The results presented in Fig 4 reinforce the fact that LdSET1 regulates the cell’s response to oxidative stress.

### LdSET1-depleted promastigotes do not display DNA damage in response to an oxidative environment

Cells in an oxidative environment experience a variety of ill-effects. To determine if the discernible differential growth patterns of LdSET1-depleted cells in response to H_2_O_2_ were due to faster recovery of these cells following exposure to the oxidative agent, or due to complete lack of response to H_2_O_2_ at concentrations upto 200 µM, we examined one of the consequences typically suffered by cells under these conditions: double strand DNA breaks (DSBs). H_2_O_2_-induced cellular oxidative stress leads to single- and double-stranded DNA breaks due to the production of hydroxyl free radicals (^•^OH) within the cell, which attack the bases and sugar groups in the double helix. In trypanosomatids, time-course kinetics of recruitment of various repair proteins to DSBs induced by infrared radiation has revealed that DSBs are primarily repaired by the HR (homologous recombination) pathway involving Exo1, RPA and RAD51 [18]. RAD51-ssDNA filaments play a major role in homology recognition and strand invasion, thus promoting eventual strand exchange in the repair process, and DSBs trigger the activation of RAD51 expression and formation of distinct RAD51 foci at the DSBs [19–21]. In examining the effect of H_2_O_2_ on the induction of DNA damage we initially adopted the route of using RAD51 as a marker for DSB repair. For this, cultures of *set1*^-/-^ and *set1*^+/+^ cells were initiated from stationary phase cultures and H_2_O_2_ (100 µM) added when cells reached a density of ∼7-9 x10^6^ cells / ml (*set1*^-/-^ cultures being initiated three days earlier to enable simultaneous H_2_O_2_ treatment of both lines), incubation carried out for five hours, the medium replaced with fresh H_2_O_2_-free medium, and cells sampled at various time intervals thereafter for isolation of whole cell lysates. The lysates were probed for RAD51 activation using anti-RAD51 antibodies already available in the lab. As seen in Fig 5A, unlike in *set1*^+/+^ parasites where RAD51 was activated over time in response to the oxidative stress induced by H_2_O_2_, in keeping with previous results from experiments with *T.cruzi* [19], RAD51 levels did not increase at any of the sampled times in *set1*^-/-^ cells. Interestingly, we observed the expression of RAD51 in untreated cells also to be significantly lower in *set1*^-/-^compared to *set1*^+/+^ cells. The reason for this is not understood at present and needs further exploration.

**Fig 5.**
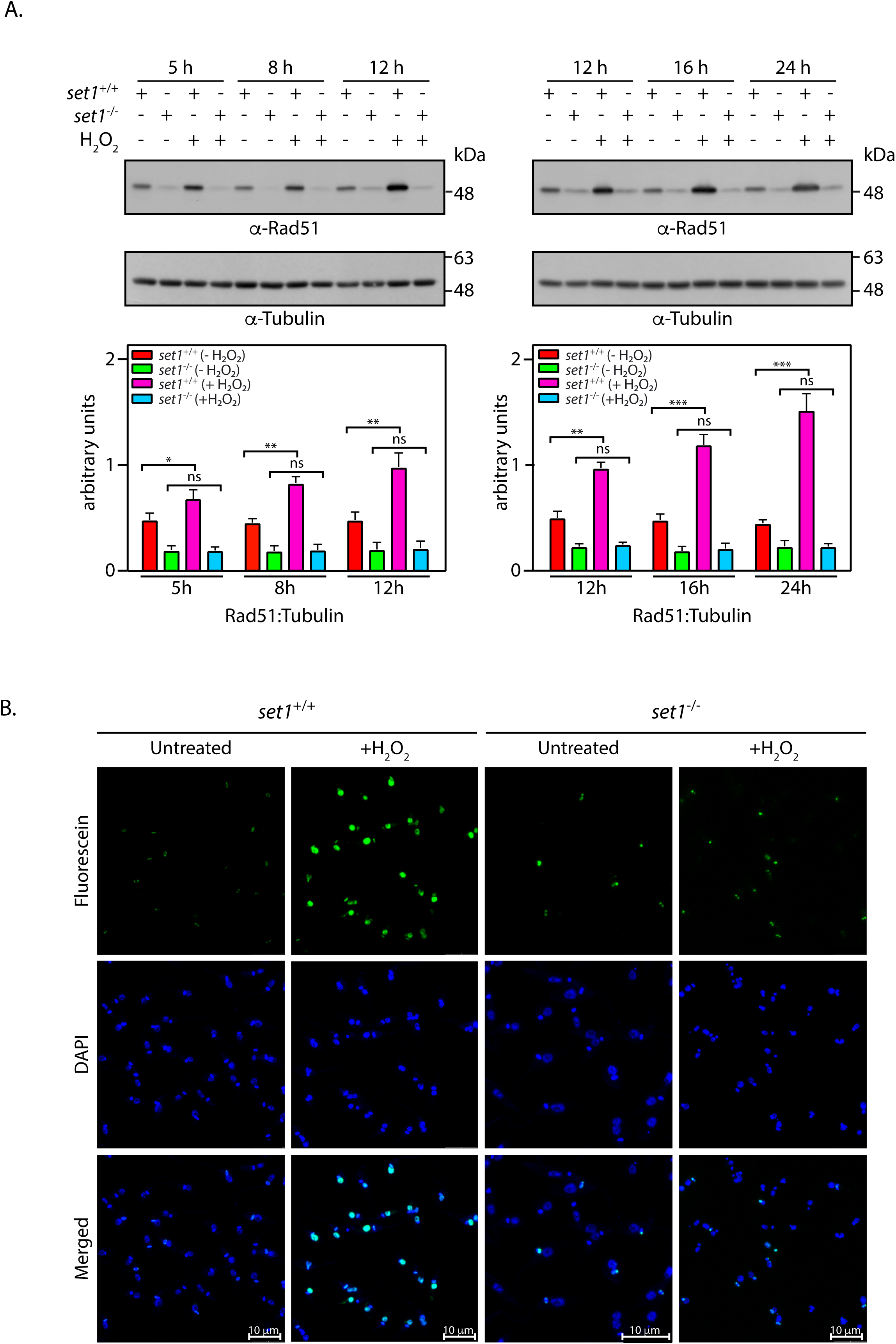
Effect of *set1* elimination on H_2_O_2_-induced DNA damage. **A.** Analysis of DNA damage response in H_2_O_2_-treated cells. Western blot analysis of whole cell lysates, isolated from 4×10^7^cells that were exposed to 100 μM H_2_O_2_ for 5 hours, using anti-RAD51 antibodies (available in the lab, 1:1000 dilution). Loading control: tubulin. Time-points indicate hours after start of the exposure. Quantitation was carried out using ImageJ analysis and plotted values represent average of three experiments. Error bars represent standard deviation and statistical significance was determined using student’s *t*-test. ***: *p* value <0.0005, **: *p* value <0.005, *: *p* value <0.05, ns: statistically not significant. **B.** Analysis of DNA damage in H_2_O_2_-treated cells. Microscopic analysis of TUNEL assay reactions that were performed on cells that were exposed to 200 μM H_2_O_2_ for 5 hours. DAPI stains nucleus and kinetoplast in each cell. Fluorescein labels free 3’OH ends of DNA, as generated by breaks. Magnification bar: 10 μm. Images were captured by Z-stack analysis using confocal microscopy. *set1^+/+^*cells: Ld1S::neo-hyg cells. *set1^-/^*^-^ cells: *set1*-nulls.

To rule out the possibility of absence of RAD51 activation in *set1*-nulls reflecting DNA repair occurring through a RAD51-independent pathway in these cells, we directly assessed DNA damage using the TUNEL assay, which detects DNA strand breaks by the TdT-mediated uptake of fluorescein-tagged dUMP at the free 3’OH groups generated by the breaks. Accordingly, *L. donovani* promastigotes (*set1*^+/+^ and *set1*^-/-^) were treated with H_2_O_2_ following the same regimen, for 5 hours, followed by analyses for DNA breaks using the TUNEL reaction. As seen in Fig 5B, untreated parasites of both types (*set1*^+/+^ and *set1*^-/-^) showed labelling of kinetoplast DNA in some cells, signifying dUMP incorporation in replicating kinetoplasts [22]. While hardly any nuclei were labelled in these cells, it was observed that almost twice as many *set1*^-/-^ cells exhibited nuclear labelling relative to *set1*^+/+^cells (S1 Table), suggesting that a higher fraction of cells might enter the apoptotic pathway in case of LdSET1-depleted cells. This possibility needs further study. Contrasting to untreated cells, *set1*^+/+^ cells displayed compelling evidence of breaks in nuclear DNA after treatment with H_2_O_2_, with more than 95% of the nuclei being strongly labelled with dUMP upon exposure to 200 µM H_2_O_2_. *set1*^-/-^ cells, however, showed almost no evidence of damage in nuclear DNA in response to similar H_2_O_2_ treatment (Fig 5B, S7 Fig and S1 Table). Taken together, the data in Fig 5 underscore the fact that *set1*^-/-^ cells do not suffer any discernible damage to nuclear DNA in response to H_2_O_2_-induced oxidative environment.

### Ectopic expression of LdSET1 in set1-nulls rescues the aberrant phenotypes associated with LdSET1 depletion

To verify that the observed phenotypes were due to LdSET1 depletion, LdSET1-FLAG was ectopically expressed in *set1*^-/-^ promastigotes as described in Methods (Fig 6A) and growth of *set1*^-/-^::SET1^+^ promastigotes monitored as earlier. Ectopic expression of LdSET1 largely rescued the growth defects of *set1-*nulls (Fig 6B). At ∼12 hours, the generation time of *set1*^-/-^::SET1^+^ promastigotes was found to be near to that of *set1*^+/+^ cells (Fig 6C). Flow cytometry analysis of hydroxyurea-synchronized promastigotes revealed an alleviation of the defects observed in *set1*^-/-^ cells, upon ectopic expression of LdSET1 in them (Fig 6D). The effect of H_2_O_2_-induced oxidative stress on *set1*^-/-^::SET1^+^ cells was examined by incubating cells with H_2_O_2_ (100 µM) and monitoring growth. The data in Fig 6E demonstrate that LdSET1-FLAG expression in *set1*-nulls allowed the parasite to largely overcome the mutant phenotype, with *set1*^-/-^::SET1^+^ promastigotes being vulnerable to H_2_O_2_ exposure almost as much as *set1*^+/+^ cells. This was also reflected in the RAD51 activation profile of *set1*^-/-^::SET1^+^ cells in response to H_2_O_2_ exposure (Fig. 6F). The observation of only a partial rescue of *set1*^-/-^ phenotypes could perhaps be due to differential expression of LdSET1-FLAG in the many cells of the population.

**Fig 6:**
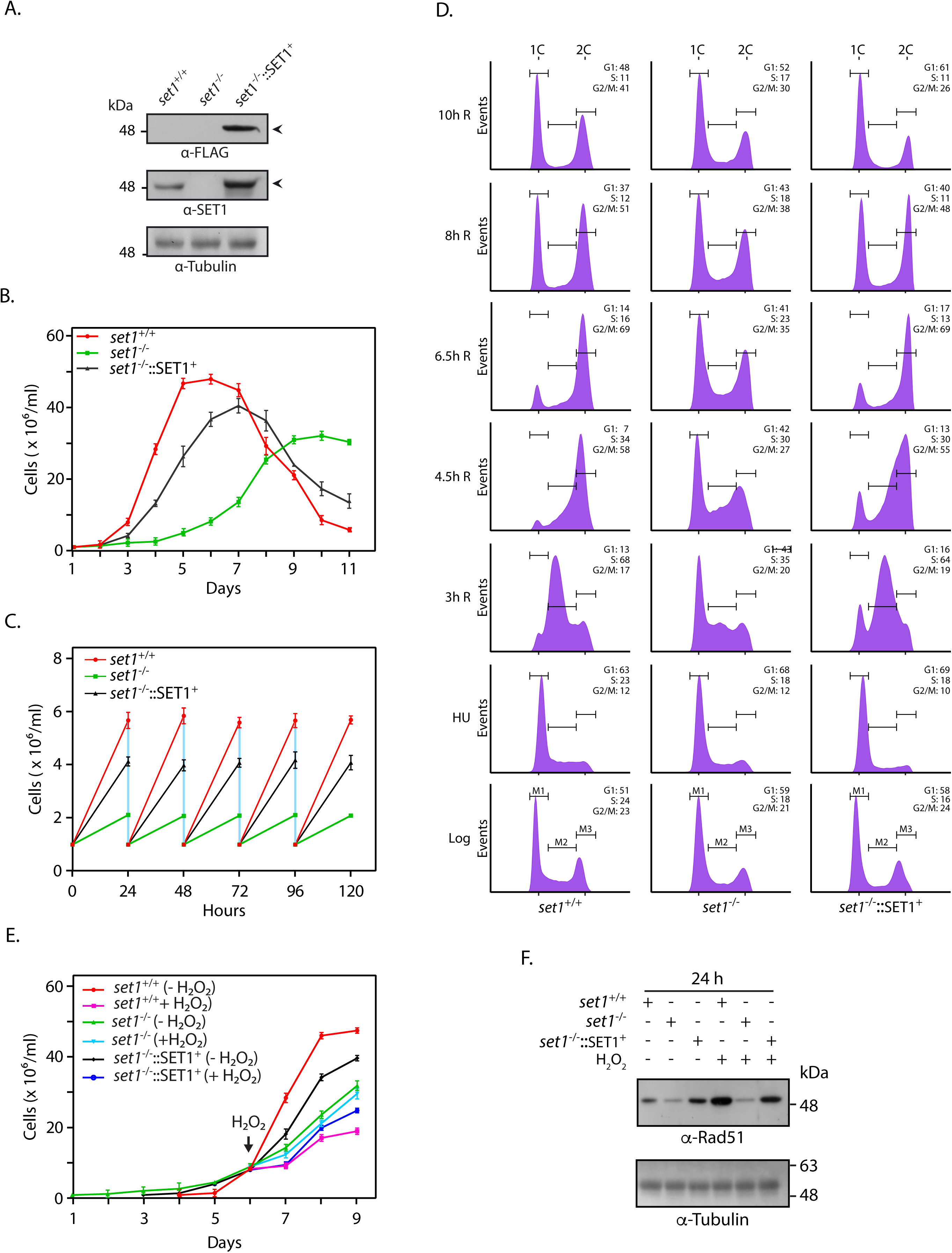
Effect of ectopic expression of LdSET1 in *set1*-nulls, on the phenotypes associated with *set1* deletion. **A.** Western blot analysis of whole cell extracts isolated from 8×10^7^ logarithmically growing promastigotes using anti-SET1 and anti-FLAG antibodies (1:1000 dilution). Loading control: tubulin. **B.** Analysis of growth of parasites. Cultures were initiated at 1×10^6^ cells/ml, from stationary phase cultures. Values plotted represent average of three experiments, with each experiment being performed with two technical replicates. Error bars represent standard deviation. **C.** Generation time, determined by initiating cultures from logarithmically growing cultures at 1×10^6^ cells/ml, and diluting the cultures to 1×10^6^ cells/ml every 24 hours after counting them. Values plotted are average of three experiments, with error bars depicting standard deviation. **D.** Flow cytometry analysis of HU-synchronized promastigotes. Sampling time-points are indicated on the left of each row of histograms. 30000 events were analyzed at every time-point. M1, M2 and M3 represent gating for cells in G1, S and G2/M respectively. Percent cells in each cell cycle phase are indicated in upper right-hand corner of the histograms. The experiment was carried out twice, with comparable results, and one data set is show here. **E.** Effect of prolonged H_2_O_2_ exposure on promastigote growth. Cultures initiated from stationary phase cultures were treated with 100 μM H_2_O_2_, for 3 days, with cells being counted every 24 hours. **F.** Analysis of DNA damage response. Western blot analysis of whole cell lysates, isolated from 4×10^7^ cells that were exposed to 100 μM H_2_O_2_ for 5 hours, using anti-RAD51 antibodies (1:1000 dilution). Loading control: tubulin. Lysates were isolated 24 hours after H_2_O_2_ exposure. *set1^+/+^*cells: Ld1S::neo-hyg cells carrying empty vector. *set1^-/^*^-^ cells: *set1*-nulls. *set1^-/^*^-^::SET1^+^ cells: Transfectant *set1*-nulls expressing SET1-FLAG ectopically.

### set1-null parasites do not exhibit activation of reactive oxygen species in response to exposure to hydrogen peroxide

The data in Figs. 2-6 strongly indicate that LdSET1 depletion modulates the cell’s response to an oxidizing environment. Particularly intriguing in *set1*-nulls was the apparent absence of detectable DNA damage, expected to be induced by the production of reactive oxygen species (ROS) upon exposure to H_2_O_2_ (Fig. 5). The production of reactive oxygen species (ROS) in *set1^-/-^* cells was compared with that in *set1^+/+^* cells using the DCFDA assay [23]. This assay is based on the principle that dichlorodihydrofluorescein diacetate (DCFDA) taken up by cells is deacetylated by cellular esterases to dichlorodihydrofluorescein. Intracellular ROS oxidize the dichlorodihydro-fluorescein to dichlorofluorescein, whose fluorescence is measured spectrofluorimetrically (498 ex/529 em). The assay was carried out by incubating *set1*^+/+^ and *set1*^-/-^ promastigotes in medium carrying H_2_O_2_ (100 µM), before incubation with DCFDA and analysis by measurement of fluorescence emission at 529 nm (as detailed in Methods). It was observed that while ROS were activated in *set1*^+/+^ cells over 45 min-24 h after exposure to H_2_O_2_ before gradually dropping, no appreciable change in ROS (in comparison with untreated cells) was detectable in *set1*^-/-^ cells over the same period of time except a slight elevation detected just after exposure to H_2_O_2_ (Fig 7A). Ectopic expression of LdSET1 in *set1*^-/-^ parasites partially rescued the mutant phenotype (Fig 7B). The apparent absence of induction of ROS production in response to H_2_O_2_ treatment in *set1*^-/-^ cells, may be the likely reason for these cells not suffering DNA damage under these conditions, and could also explain why the parasites are apparently resistant to fairly high concentrations of H_2_O_2._ This observation also pointed toward the possibility of a significantly higher efficiency of ROS scavenging in *set1*^-/-^ parasites.

**Fig 7.**
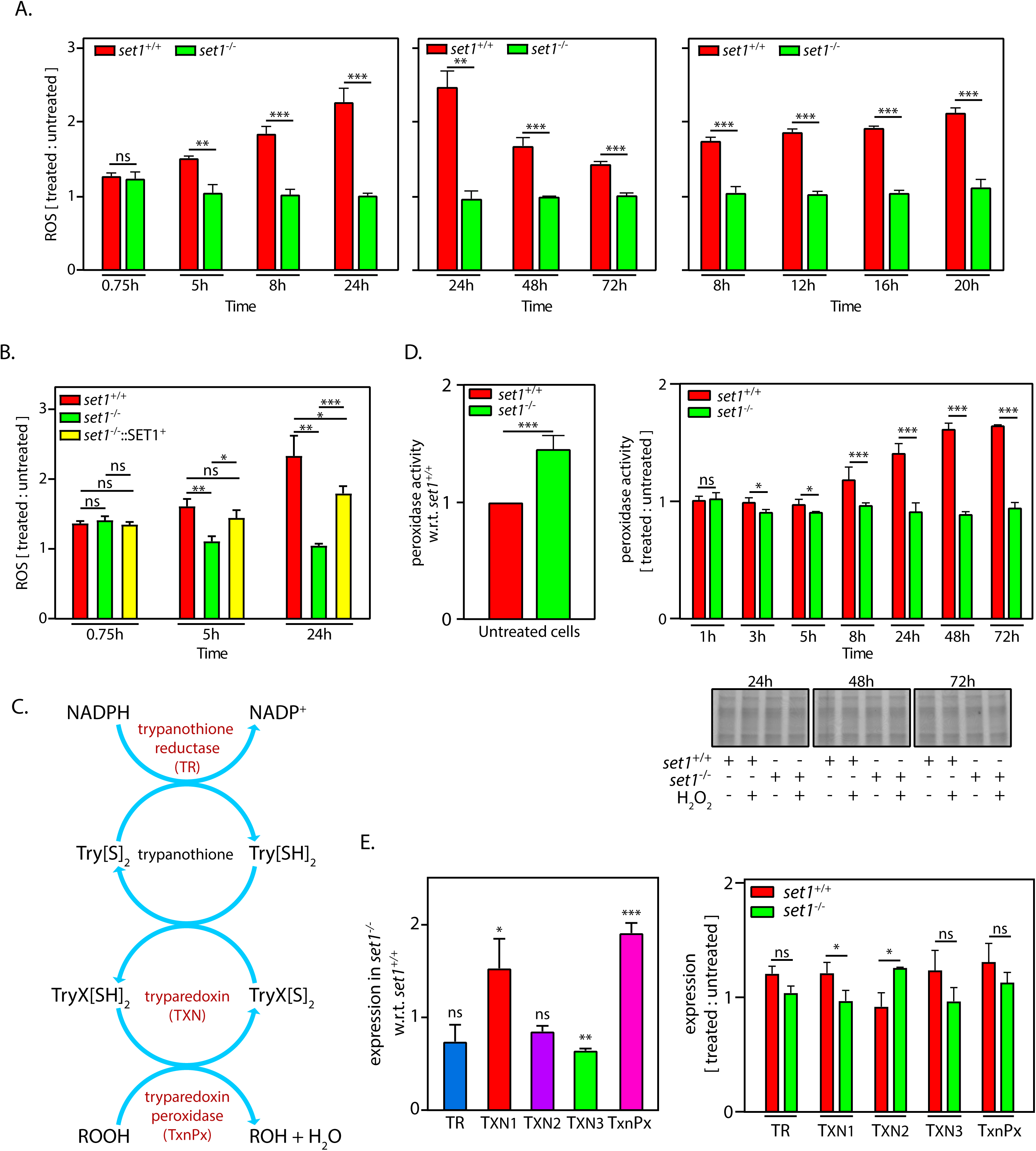
Effect of *set1* deletion on ROS production. **A.** Analysis of ROS production in *set1^+/+^*and *set1^-/-^* cells treated with 100 μM H_2_O_2_, relative to the respective cells which had not been exposed to H_2_O_2._ Sampling time is marked with reference to time of start of H_2_O_2_ treatment. The three graphs represent three separate experiments that were each performed thrice (as detailed in Methods), and values plotted are average of three experiments, with error bars depicting standard deviation. Statistical significance was determined using student’s *t*-test. ***: *p* value <0.0005, **: *p* value <0.005, ns: statistically not significant. **B.** Analysis of ROS production in *set1^+/+^*, *set1^-/-^* and *set1^-/^*^-^::SET1^+^ cells treated with 100 μM H_2_O_2_, relative to respective cells which had not been exposed to H_2_O_2._ Values plotted are average of three experiments, with error bars depicting standard deviation. Statistical significance was determined using student’s *t*-test. ***: *p* value <0.0005, **: *p* value <0.005, *: *p* value <0.05, ns: statistically not significant. **C.** Schematic representation of trypanothione peroxidase scavenging system in trypanosomatids. **D.** Analysis of peroxidase activity (as detailed in Methods). Left panel: activity in untreated *set1^-/-^* cells relative to untreated *set1^+/+^* cells. Right upper panel: activity in *set1^+/+^* and *set1^-/-^* cells treated with 100 μM H_2_O_2_, relative to respective cells which had not been exposed to H_2_O_2._ Sampling times are with reference to start of H_2_O_2_ treatment. Right lower panels: Coomassie-stained gels of cell inputs used in the reactions (input loading controls). The experiment was performed thrice (as detailed in Methods), and values plotted are average of three experiments, with error bars depicting standard deviation. Statistical significance was determined using student’s *t*-test. ***: *p* value <0.0005, *: *p* value <0.05, ns: statistically not significant. **E.** Real time PCR analysis of transcripts of the trypanothione peroxidase system. Fold difference in expression was determined using 2^-△△^_Ct_ method. Tubulin served as internal control for each RNA sample. Left panel: expression in untreated *set1^-/-^* cells with reference to untreated *set1^+/+^* cells. Right panel: expression in H_2_O_2_- treated *set1^+/+^* and *set1^-/-^* cells with reference to the respective untreated cells. TR: trypanothione reductase; TXN1, TXN2 and TXN3: tryparedoxin 1, tryparedoxin 2, and tryparedoxin 3; TxnPx: tryparedoxin peroxidase. Data plotted are average of three experiments, each experiment being performed with technical duplicates. Error bars signify standard deviation. Statistical significance was assessed using student’s *t*-test. ***: *p* value <0.0005, **: *p* value <0.005, *: *p* value <0.05, ns: statistically not significant.

Whereas the mammalian defense against oxidative stress is largely glutathione-dependent, with intracellular superoxide radicals being converted to molecular oxygen and hydrogen peroxide by superoxide dismutase (SOD), and hydrogen peroxide in turn being converted to water and oxygen by enzymes like glutathione peroxidase and catalase, trypanosomatids employ a somewhat different modus operandi. Lacking catalase and selenium-dependent peroxidases, instead, they possess a unique trypanothione-dependent system with a set of enzymes that act concertedly for the detoxification of peroxides. The three main components of this system are trypanothione reductase (TR), tryparedoxin (TXN) and tryparedoxin peroxidase (TxnPx) (Fig 7C). While TR reduces the thiol trypanothione (TS_2_) to T(SH)_2_ using NADPH, TXN (a thiol disulphide oxidoreductase) serves as the conduit through which the reducing equivalents flow from T(SH)_2_ to TxnPx (a 2-Cys peroxiredoxin), and the reduced TxnPx reacts with and catalyzes the breakdown/reduction of hydroperoxides. The absence of detectable ROS induction in response to H_2_O_2_ treatment led us to consider if this trypanothione-dependent system was overactive in *set1^-/-^* cells. Thus, we examined peroxidase activity in *set1^-/-^* parasites, as described in Methods. Results of activity assays revealed that untreated *set1^-/-^* parasites possessed significantly higher peroxidase activity relative to untreated *set1^+/+^* parasites (Fig 7D, left panel). Interestingly, while peroxidase activity was enhanced in *set1^+/+^* parasites over 45 min to 72 hours after exposure to H_2_O_2_, *set1^-/-^* parasites did not demonstrate any visible change in peroxidase activity in response to H_2_O_2_ over the same period, suggesting that elimination of LdSET1 was causing loss of regulation of peroxidase activity, and the higher levels of basal peroxidase activity in *set1^-/-^* cells were sufficient to rapidly scavenge ROS (Fig 7D).

In contemplating the reasons for enhanced basal peroxidase activity in *set1*-nulls, the possibility of higher expression of tryparedoxin peroxidase was considered. We were unable to check protein expression levels directly in the absence of available antibodies. Thus, we analyzed transcript levels of the enzyme, as well as of the other components of the trypanothione-dependent peroxide detoxification system, in *set1^-/-^* cells in comparison with *set1*^+/+^ cells. This was done by real-time PCR analysis of RNA isolated from untreated and H_2_O_2_-treated (8 h after start of the 5-hour treatment with 100 µM H_2_O_2_) cells. On comparing expression in *set1^-/-^* and *set1*^+/+^ untreated cells, it was observed that while TR was expressed poorer in *set1^-/-^* cells, TxnPx was expressed almost two-fold higher in these cells (Fig 7E, left panel). The poorer expression of TR may be compensated for by the higher expression of one of the three TXN genes (TXN1). In response to H_2_O_2_ treatment, none of the transcripts were significantly elevated in either type of parasite (Fig 7E, right panel). The enhanced peroxidase activity in *set1^+/+^*cells in response to H_2_O_2_ exposure reflects previous results from Iyer *et al*., who reported activated expression of the tryparedoxin peroxidase protein in *L. donovani* in response to H_2_O_2_ treatment [24]; our data indicates that this is not due to elevated transcripts.

The data in Figs 7D-E suggest that LdSET1 regulates peroxidase activity by controlling TxnPx expression: deletion of *set1* is coupled to significantly higher levels of *txnPx* transcripts in the cells regardless of exposure to an oxidative environment, in turn resulting in higher peroxidase activity which probably leads to rapid scavenging of the ROS produced in the cell in response to H_2_O_2_ treatment, such that ROS activation escapes detection and the effects of ROS activation (such as DNA damage) are not experienced.

## Discussion

As the cause of Visceral Leishmaniasis, the mechanisms by which *L. donovani*’s physiological processes are regulated remain an area of intensive research. Gene regulation is somewhat unusual in *L. donovani* and other trypanosomatids, with the genes being organized in long unidirectional clusters of functionally unrelated genes that are by-and-large coordinately transcribed from divergent strand switch regions (dSSRs). No consensus sequences or defined promoter elements have been recognized at these Transcription Start Regions (TSRs), nor have the wide repertoire of canonical transcriptional activators found in more conventional eukaryotes, been identified as yet in these organisms. Epigenetic mechanisms appear to control gene expression in trypanosomatids to a fair extent, and the roles of specific histone acetylations in modulating transcription and other DNA-related processes in *Leishmania* as well as *Trypanosoma* species have been documented [25–31]. The functions of histone methylations and the proteins mediating them, however, remain largely unexplored in these organisms, with studies thus far revealing the enrichment of H3K4 methylation marks at TSRs [31, 32], and the role of the *Trypanosoma* Dot1 proteins in modulation of DNA replication (via H3K76 methylation) being uncovered [33]. Histone methylations at lysine residues are mediated by SET-domain proteins (Dot1 proteins being the only other histone lysine methyltransferases), and while 29 SET-domain proteins have been identified in *T. brucei*, all of which are conserved in *L. donovani*, their target substrates have not been identified yet, and the cellular role of only one of them (TbSET27) has been reported thus far [12, 13]. SET proteins that have been extensively studied across eukaryotes, particularly yeast and mammalian cells, have been found to target non-histone substrates as well [1, 34] - highlighting the fact that they regulate cellular processes through a wide range of downstream substrates. In several instances one or more functions of a SET protein have been unearthed without the target substrate(s) being identified. These proteins often display stringent catalytic activity as they may mediate mono-, di, or tri-methylation of the target lysine residue, translating into tighter regulation of the pertinent cellular process as the extent of methylation of a particular lysine residue may govern protein-protein interactions. SET protein-mediated methylations also impact protein activity and protein stability.

In investigating the SET1 protein in *L. donovani*, we found the primarily cytosolic LdSET1 was expressed in promastigotes throughout the cell cycle (S2C and S3C Fig). LdSET1 was not essential to the parasite, but *set1^-/-^* promastigotes exhibited slower growth and a heightened sensitivity to HU-induced G1/S arrest in comparison to *set1^+/+^*cells (Fig 3). Contrastingly, *set1^-/-^* amastigotes survived and proliferated more proficiently than *set1^+/+^* amastigotes within host macrophages (Fig 4A), indicating that LdSET1 plays a role in moderating the parasite’s response to the hostile intracellular oxidative environment of host cells. Interestingly, this effect was tightly controlled by LdSET1 expression levels: partial depletion of LdSET1 to ∼50% its usual expression, while not having any effect on growth or cell cycle progression of promastigotes (Figs 1B, 1C), led to much higher proficiency of survival and propagation of amastigotes in host macrophages (Fig 1D), even higher than that of *set1^-/-^* amastigotes (Fig 4A). The impact of LdSET1 depletion on amastigote growth and survival was mirrored in the response of promastigotes to an oxidative growth environment *in vitro*, wherein partial depletion of LdSET1 allowed the parasites to tolerate H_2_O_2_-induced oxidative stress better than usual (Fig 2), while complete elimination of LdSET1 made the parasites almost completely resistant to stress induced by H_2_O_2_ at concentrations upto 200 µM (Figs 4B, 4C). The resistance of *set1^-/-^* promastigotes to H_2_O_2_-induced stress was apparent not only in the growth kinetics of the parasites, but also in the almost complete absence of DNA damage in response to H_2_O_2_ exposure (Fig 5, S7 Fig and S1 Table). ROS-induced stress in mammalian cells is known to alter the subcellular localization of the p65/RelA subunit of the NF-kB transcription protein complex, relocating the cytoplasmic protein to the nucleus where it exercises its ability to activate transcription of several genes such as Mn-SOD and catalase [34, 35]. No obvious alteration in the subcellular localization of LdSET1 was detectable in response to H_2_O_2_ exposure (S6B Fig), and LdSET1 expression was not upregulated upon H_2_O_2_ exposure either (Fig 4D). The finding that LdSET1 elimination makes the parasite resistant to H_2_O_2_-induced oxidative stress and allows amastigotes to propagate more proficiently suggests that the parasite (wild type) harmonizes its response with the host environment, to withstand the onslaught of the host defense system and proliferate in an orderly fashion, thus establishing itself firmly in the host. The higher proliferation rates associated with *set1* deletion would lead to a depletion of host cells with time, thence being detrimental to the persistence of infection.

Amastigotes express a gamut of proteins to tolerate and overcome the stress induced by the oxidative environment of the macrophages. While the macrophages assault the parasites early in infection by releasing a burst of reactive oxygen species (ROS), the parasite employs a barrage of proteins to fight the ROS wave. These include anti-oxidant proteins and enzymes to scavenge the ROS, as well as proteins to repair DNA lesions induced by the oxidative environment. Hydroperoxides in *Leishmania* are scavenged by a trypanothione-dependent system. Three tryparedoxin (TXN) genes have been annotated in the *L. donovani* genome, and of the two tryparedoxin peroxidases (TxnPx) that have been identified in *Leishmania*, one is cytosolic and essential for cell survival, while the other is mitochondrial [24, 36]. The cytosolic TxnPx is the primary mediator of hydroperoxide scavenging in the macrophage environment, and has been reported to be secreted in *L. infantum* and *L. major* [37]. The data in Fig 7 indicates that LdSET1 controls expression and activity of tryparedoxin peroxidase through transcript levels (Figs 7D, 7E), with *set1* deletion leading to rapid scavenging of any ROS generated (Fig 7A).

While the mechanism by which LdSET1 controls *txnPx* transcript levels remains unknown to us, two general possibilities are contemplatable. In the first scenario, LdSET1 may regulate *txnPx* transcripts either through an epigenetic mark on a histone residue, which would have an impact on global gene expression, or through the methylation of a transcription factor which downregulates *txnPx* transcription. Considering that LdSET1 is predominantly cytosolic, it is unlikely to mediate an activating/repressive histone methylation mark, which would usually be added after histone deposition. While *txnPx* transcripts are upregulated in *set1*-nulls (Fig 7E), other genes lying in the same polycistronic cluster remain unaffected by *set1* deletion in untreated as well as H_2_O_2_-exposed cells (data not shown), indicating that LdSET1 does not modulate global gene activation/repression. Contemplating the alternate possibility of LdSET1 controlling the expression of *txnPx* transcripts through the methylation of a transcriptional activator: previous studies have reported SET protein-mediated methylation to impact activity of such proteins. Intracellular ROS accumulation in mammalian cells shuttles the p65/RelA subunit of the NF-kB transcription factor into the nucleus, ultimately triggering activation of genes whose products fight the oxidative burst. While Set7/9-mediated methylation of p65/RelA at a specific lysine residue is believed to be critical to RelA’s ability to activate transcription, methylation of RelA at other lysine residues by the same Set protein has been reported to repress its ability to activate transcription (reviewed in [34]). The abundantly cytosolic human SET protein SMYD2 methylates p53 which shuttles between the nucleus and cytoplasm, and this methylation (at K370) inhibits p53 transactivation activity [38]. Thus, LdSET1-mediated methylation of a transcriptional activator may inhibit its activity, tightly controlling *txnPx* transcript levels, and deletion of *set1* may alleviate this negative regulatory effect (Fig 8, “1?”). While no evidence in support of gene regulation through transcriptional activators has been uncovered in trypanosomatids thus far, with transcription being primarily polycistronic and constitutive, a small subset of genes have been identified to be transcriptionally activated in a cell cycle-dependent manner. These genes are scattered throughout the genome, and are turned on by promoters immediately upstream of them [25]. It is possible that *txnPx* is activated by a promoter of this kind; this needs to be further investigated.

**Fig 8.**
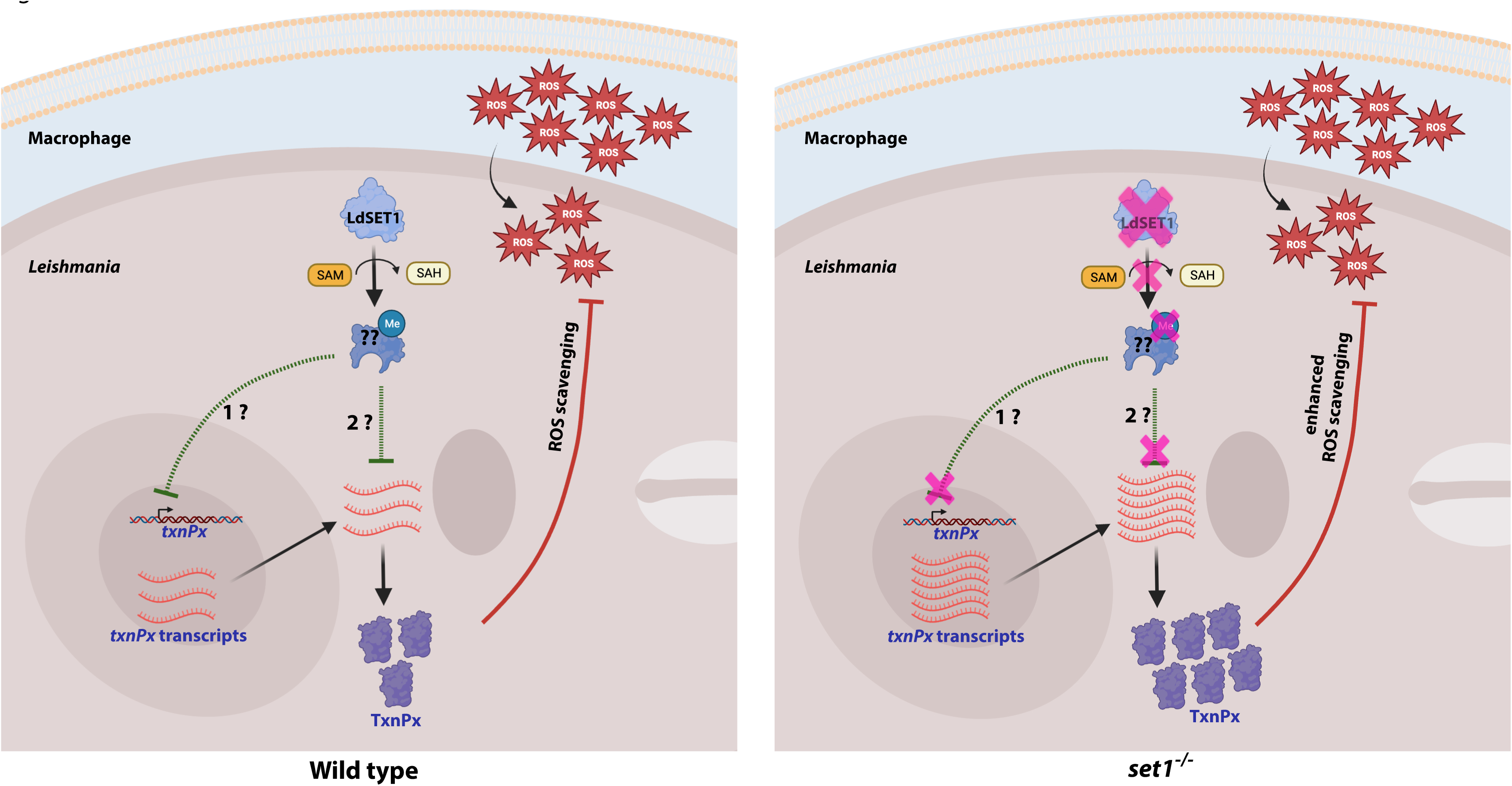
The role of LdSET1 in moderating parasite response to oxidative stress. LdSET1-mediated methylation of an unidentified target protein may either negatively impact transcriptional activation of the *txnPx* gene (labelled “1?” in the figure), or negatively impact stability of the processed transcripts (labelled “2?” in the figure). Deletion of the *set1* gene removes this negative regulatory effect, culminating in higher peroxidase activity, enhanced ROS scavenging, and thus higher tolerance to the oxidative milieu. Figure created with BioRender software (BioRender.com).

In the second scenario, in light of the fact that post-transcriptional processes are crucial to regulating gene expression in these organisms, the role of RNA-binding proteins (RNA-BPs or RBPs) may be a vital factor. 5’ and 3’ UTRs of stable processed transcripts often carry *cis*-acting elements that serve as regulatory motifs to which RBPs bind. The RBP-RNA interactions modulate mRNA stability, decay, transport, and can also modulate translation efficiency. The *T. cruzi* UBP1 protein has been reported to bind to a specific sequence UBP1m in the 3’UTR of the gene transcripts whose expression it regulates (mostly those encoding surface glycoproteins), thus stabilizing the mRNAs [39]. While PTMs on proteins have been widely studied and are generally known to regulate protein function through their impact on localization, protein-protein interactions, stability and activity, not much information is currently available on the impact of PTMs in case of RNA-BPs. An in-depth analysis of the PTM landscape on RBPs in human cells, has been carried out through collation of existing curated datasets and published research [40]. PTMs have been identified in almost 2400 RBPs, with the vast majority of these carrying multiple PTMs. The commonest classes of PTMs identified were phosphorylation, acetylation, ubiquitination and methylation. Methylations on RBPs have been found in over 1100 proteins and while typically occurring at arginine residues, methylation of RBPs at lysine residues have also been identified in several human cancer cell lines as well as in clinical samples drawn from different body organs [40]. Thus, it is possible that LdSET1 mediates its effect through the methylation of one or more RBPs, the negative impact of which would be reflected in tightly controlled levels of target transcripts – in this case, *txnPx.* Depletion of LdSET1 and corresponding abrogation of the RBP methylation event would be coupled to loss of this regulation, resulting in an overall increase in *txnPx* transcript expression levels (Fig 8, “2?”). The mammalian SET7/9 - mediated E2F1 methylation marks it for degradation via the ubiquitination-proteasomal degradation pathway [41]; a similar mechanism could gradually deplete the parasite of an RNA-BP that is vital to stabilizing *txnPx* transcripts.

This study reports the first data directly investigating the functional role of a SET protein in *Leishmania* species, with the results of our experiments demonstrating that LdSET1 modulates the persistence of *Leishmania* infection by working towards maintaining the delicate balance between parasite and host cell. The distinct phenotypes obtained upon *set1* deletion suggest that although 29 SET proteins have been identified in these organisms, functional redundancy may be limited. While the global landscape of histone modifications has not been uncovered yet in *Leishmania* species, a large number of methylation events have been identified in *T. brucei* and *T. cruzi* histones, and several of these SET proteins must target histone substrates. The identification of target histone substrates of specific SET proteins will allow us to gain new insights into how these proteins function.

## Materials and Methods

### Leishmania cultures and manipulations

*L. donovani* 1S (Ld1S) cultures were grown in M199 medium (Lonza, Switzerland) supplemented with fetal bovine serum (Invitrogen, USA), adenine and hemin (Sigma Aldrich, USA), at 26°C as described earlier [42]. Growth and survival patterns were analyzed as described earlier [27]. Generation time in the logarithmic phase of growth was determined as described earlier [25]. Synchronization of parasites using hydroxyurea (5 mM for 8 hours) and flow cytometry analyses were carried out as described previously [17]. Transfections were done and clonal lines generated as described [26, 27]. Whole cell extracts were isolated using the M-PER kit (Thermo Fisher Scientific, USA) as per manufacturer’s instructions. Procylics and metacyclics were isolated as described earlier [43]. For treatment with hydrogen peroxide, cultures were initiated from stationary phase cultures, at 1×10^6^ cells/ml, and hydrogen peroxide (25-1000 μM) added on reaching a cell density of ∼ 7-9 x10^6^ cells/ml (Day 3 for *set1*^+/+^ and *set1* ^-/+^ cells, Day 6 for *set1*^-/-^ cells). The *set1*^-/-^ cultures were initiated three days earlier to enable addition of hydrogen peroxide at the same time as the *set1^+/+^* and *set1* ^-/+^ cells. At the time of addition of H_2_O_2_ the cultures were split into two, with half the cells receiving H_2_O_2_ treatment and the other half being carried forward as the untreated culture. The cultures were incubated further at 26°C for varying time intervals, and sampled for further experimentation.

### Cloning of set1 gene for expression in Leishmania

For expression in *Leishmania* parasites, the pLEXSY-FLAG vector was created from the pLEXSY-CYC9-FLAG plasmid [25] by digesting it with BglII (which would drop out the CYC9 gene but not the FLAG sequence it is tagged with), and ligating the vector’s BglII ends. For expressing the *set1* gene in *L. donovani* promastigotes, the gene was amplified off *L*. *donovani* 1S genomic DNA using primers SET1-FLAG-F (5’-TAGGATCCATGCCCATCAGCCAG-3’) and SET1-FLAG-R (5’-TAGGATCCAGGAAGAAGAGGCTTCA-3’). The amplicon was cloned into the BglII site of the pLEXSY-FLAG vector using the BamHI sites in the primers, creating plasmid pLEXSY/SET1-FLAG.

### Creation of knockout and rescue lines

The first *set1* genomic allele was knocked out by replacing it with a *hyg^r^* cassette, while the second allele was replaced with a *neo^r^* cassette. The donor plasmid for replacing the second allele was constructed using the pLEXSY_I-neo3 vector as backbone (Jena Bioscience, Germany). To replace the first allele, first the *neo^r^*cassette in pLEXSY_I-neo3 was replaced with a *hyg^r^* cassette using the BamHI-SpeI sites flanking the *neo^r^* cassette, generating the vector pLEXSY/hyg, which served as the backbone for constructing the donor plasmid. The ∼800 bp region immediately upstream of the *set1* gene was amplified off Ld1S genomic DNA using primers SET1-5’FL-F (5’-TAGCGGCCGCATTTAAATGGTTTTTTCTGCCTTCTCTTG-3’) and SET1-5’FL-R (5’-TAGCGGCCGCGGTCTGCGACCGATATACCTCGG-3’), and the amplicon cloned into the NotI site of the two vector backbones. The ∼ 800 bp region immediately downstream of the *set1* gene was then amplified using primers SET1-3’FL-F (5’-TAACTAGTATACATGAGTGAGACGCTCC GCGG-3’) and SET1-3’FL-R (5’-TAACTAGTATTTAAATTGCTTCTCAAAGCCCTTGTCA-3’), and the amplicon cloned into the SpeI site of the two clones carrying the 5’flank sequence - thus creating the donor plasmids SET1-KO/hyg and SET1-KO/neo.

For making the *set1* heterozygous knockout (*set1*^-/+^) the donor cassette was released from plasmid SET1-KO/hyg using SwaI digestion, transfected into *L. donovani* promastigotes by electroporation, and clonal lines selected for and expanded in the presence of hygromycin (16 μg/ml), as described earlier [26]. For making the *set1*-null (*set1*^-/-^) the donor cassette was released from plasmid SET1-KO/neo using SwaI digestion, transfected into *set1*^-/+^ promastigotes by electroporation, and clonal lines selected for and expanded in the presence of G418 (50 μg/ml) and hygromycin (16 μg/ml). Clonal lines were maintained in liquid culture under selection pressure (G418 at 100 μg/ml and hygromycin at 32 μg/ml) except for flow cytometry experiments where the drugs were withdrawn a week before setting up the experiment.

For making the rescue line, the *set1* gene was cloned into the BamHI-EcoRV sites of pXG-FLAG (bleo) vector [27], using primers SET1-pXG-F (5’-GAGGATCCGC CACCATGCCCATCAGCCAG-3’) and SET1-pXG-R (5’-AGGATATCTCCAGGAAGAAGAGGCTT-3’) to amplify the gene for cloning. The plasmid pXG-SET1-FLAG (bleo) was transfected into *set1*^-/-^ promastigotes and clonals selected for using G418, hygromycin and phleomycin (50 μg/ml, 16 μg/ml and 2.5 μg/ml respectively).

### Isolation of RNA and real time PCRs

Total RNA was isolated from 5×10^7^ promastigotes with the help of the PureLink RNA mini kit (Invitrogen, USA). The RNA was treated with DNaseI (1U DNase I per 2 μg RNA at 37°C for 30 min) prior to cDNA synthesis, to eliminate any genomic DNA contamination. Total cDNA was synthesized as per the manufacturer’s instructions using the iScript cDNA synthesis kit (Bio-Rad, USA). For expression analysis using real-time PCR, one-twentieth of the cDNA synthesis reaction was used as template per reaction. Tubulin expression was analyzed using primers Tubulin RT-F1 (5’-CTTCAAGTGCGGCATCAACTA-3’) and Tubulin RT-R2 (5’-TTAGTACTCCTCGACGTCCTC-3’). Trypanothione reductase expression was analyzed using primers designed against LdBPK_050350.1 (TR-RT-F: 5’- CACAACATCAGCGGCAGCAAG-3’ and TR-RT-R: 5’-TCGGCGCTCGTCGGGTGGA-3’). Expression of Tryparedoxin 1 was analyzed using primers designed against LdBPK_291250.1 (TXN1-RT-F: 5’- GAGTTCTACGAGAAGCATCACA-3’ and TXN1-RT-R: 5’-TCAGCGTCGGAATCGATTCCA-3’). Expression of Tryparedoxin 2 was analyzed using primers designed against LdBPK_291240.1 (TXN2-RT-F: 5’- CAACAAACA CGCGAAGTCGAAG-3’ and TXN2-RT-R: 5’-CGACGCCGATCAGCGTCGGA-3’). Expression of Tryparedoxin 3 was analyzed using primers designed against LdBPK_312000.1 (TXN3-RT-F: 5’- GACTACTACTGCCTGCCGTAC-3’ and TXN3-RT-R: 5’- GGCTGCTGC GGCTCTGCATC-3’). Expression of Tryparedoxin peroxidase was analyzed using primers designed against LdBPK_151140.1 (TxnPx-RT-F: 5’- GCCTACCGCGGTCTCTTCATC-3’ and TxnPx-RT-R: 5’-TTCCAGTTCGCGGGGCACAC-3’). Expression of genes in *set1*^-/-^ cells relative to in *set1*^+/+^ cells were determined using the 2^-△△^_Ct_ method [44], with tubulin gene expression serving as the internal control in each sample-type. Expression of genes in H_2_O_2_-treated versus untreated cells were likewise determined using the 2^-△△^_Ct_ method using the tubulin gene as internal control. Real time PCR experiments were done three times, with technical duplicates in each experiment. Values plotted are the average of three experiments, and error bars show standard deviation. Student’s *t-test* was applied for analyzing statistical significance.

### Immunofluorescence analysis

Indirect immunofluorescence was carried out as described earlier [27]. Briefly, exponentially growing promastigotes expressing LdSET1-FLAG were fixed with 2% paraformaldehyde, cell spreads prepared on poly-lysine coated coverslips, cells permeabilized with 0.1% Triton-X100, blocked with 5% chicken serum, incubated with anti-FLAG antibody for two hours (Sigma Aldrich, 1:100 dilution), antibody washed off, incubated with Texas Red-labelled secondary antibody for an hour (Jackson Immunoresearch Laboratories, USA, 1:100 dilution), and finally mounted in anti-fade solution containing DAPI (Vectashield, Vector Laboratories, USA). Cells were viewed using the Leica TCS SP8X confocal microscope (with 100X (in oil) objective), and images captured and analyzed using the Leica Application Suite X software.

### TUNEL assay

To assess DNA strand breaks in *L. donovani* promastigotes (*set1*^+/+^ and *set1*^-/-^), cultures initiated at 1×10^6^ cells/ml from stationary phase cultures were grown to a cell density of 7-9 x10^6^ cells/ml (Day 3 in case of *set1*^+/+^ and Day 6 in case of *set1*^-/-^ cells; *set1*^-/-^ cultures were initiated three days earlier to enable addition of hydrogen peroxide to all cultures at the same time), and hydrogen peroxide (100 μM or 200 μM) added. Cultures were incubated further for 5 hours, and aliquots of 1×10^7^ cells removed immediately after to perform the TUNEL assay. For this, the cells were collected by centrifugation at 1448*g*, washed with 1X PBS, fixed with 2% paraformaldehyde, and cell spreads prepared on poly-lysine coated coverslips (2-4 x10^6^ cells per coverslip). After cell permeabilization with 1X PBS-0.2% Triton X-100 for 10 min, the TUNEL assay was performed using the DeadEnd Fluorometric TUNEL System (Promega, USA) as per the manufacturer’s instructions. Briefly, the adhered cells were incubated with the dUTP tailing reaction mix at 37°C for an hour before stopping the reaction, washing the coverslips with 1XPBS to remove unincorporated dUTP, and mounting in anti-fade solution carrying DAPI. Cells were viewed using the Leica TCS SP8X confocal microscope (with 100X (in oil) objective), and images captured and analyzed using the Leica Application Suite X software.

### Measurement of ROS

ROS in *L. donovani* promastigotes were assessed using the DCFDA (dichlorodihydrofluorescein diacetate) assay as described [23]. Towards this, *L. donovani* promastigote cultures (*set1*^+/+^ and *set1*^-/-^) were initiated at 1×10^6^ cells/ml from stationary phase cultures, and hydrogen peroxide (100 μM) added when cells reached a density of 7-9 x10^6^ cells/ml (Day 3 in case of *set1*^+/+^ and Day 6 in case of *set1*^-/-^; *set1*^-/-^ cultures were initiated three days earlier to enable addition of hydrogen peroxide to all cultures at the same time). Cultures were incubated further at 26°C for 45 minutes, the medium replaced with fresh H_2_O_2_-free medium, and incubation at 26°C continued. Aliquots of 1×10^7^ cells were removed at various time intervals thereafter to perform the DCFDA assay.

For this, the cells were collected by centrifugation at 1448*g*, medium completely aspirated, and collected cells washed twice in HEPES/NaCl buffer (21 mM HEPES (pH 7.0), 137 mM NaCl, 5 mM KCl, 6 mM glucose, 0.7 mM NaH_2_PO_4_) before suspending the cells in 1 ml of the same buffer. The DCFDA reagent (5 μM; Sigma Aldrich) was added to the cell suspension, reaction mixed by inverting the tube, and incubated in the dark at 26°C for 45 minutes. This was followed by collecting the cells by centrifugation, washing them in the HEPES/NaCl buffer, and making a cell suspension in 1 ml of the buffer. 200 μl aliquots of the cell suspensions were excited at 488 nm and fluorescence emission detected at 529 nm using a Tecan plate reader (Infinite 200 PRO).

Control reactions carried out without addition of DCFDA yielded no fluorescence. Control reactions carried out without cells gave “background” fluorescence readings, and the values of these control reactions (which were set up at every time-point) were subtracted from the values obtained in the reactions with the two cell types (*set1^+/+^* and *set1 ^-/-^*). At every time-point, reactions with the two cell types were analyzed in case of both, untreated and treated cells. The ratio between values obtained (after subtracting control reaction value) under treated versus untreated conditions, was used as a measure of ROS activation in response to H_2_O_2_, for each cell type. The experiments were done thrice and mean values are presented in the bar charts; error bars indicate standard deviation. Student’s *t-test* was used to analyze statistical significance.

### Measurement of peroxidase activity

To analyze peroxidase activity in *L. donovani set1*^+/+^ and *set1*^-/-^ promastigotes, cultures were initiated at 1×10^6^ cells/ml from stationary phase cultures, and incubated to a cell density of 7-9 x10^6^ cells/ml (*set1*^-/-^ cultures were initiated three days earlier so that it reached the cell density at the same time as *set1*^+/+^ cells), before adding hydrogen peroxide (100 μM) and allowing incubation at 26°C for 5 hours, then replacing the medium with fresh H_2_O_2_-free M199 and continuing incubation at 26°C. Aliquots of 1×10^7^ cells were removed at various time intervals thereafter to perform the Amplex Red assay [45, 46]. For this, the cell aliquots were washed with 1X PBS and resuspended in 500 μl assay mix (1X PBS carrying 64 μM digitonin, 10 μM Amplex Red (Invitrogen), 1 mM H_2_O_2,_ 1 mM protease inhibitors mix). The reaction mixes were incubated in the dark at 26°C for 30 min, cell remnants removed by centrifugation, and fluorescence of the supernatant (200 μl aliquot) measured (excitation at 535 nm, emission at 590 nm).

Reactions carried out without addition of Amplex Red yielded no fluorescence. Reactions carried out without cells gave “background” fluorescence readings, and the values of these control reactions (set up with every time-point) were deducted from the values obtained in the reactions with the two cell types (*set1^+/+^* and *set1 ^-/-^*). At every time-point, reactions with the two cell types were analyzed in case of both, untreated and treated cells. The ratio between values obtained (after subtracting control reaction value) under treated versus untreated conditions, was used as a measure of activation of peroxidase activity in response to H_2_O_2,_ for each cell type. The ratio between values obtained in *set1*-nulls versus *set1^+/+^* in cells that had not been treated with H_2_O_2,_ was used as a measure of basal peroxidase activity in *set1*-nulls relative to *set1^+/+^* cells. The experiments were done thrice and mean values are presented in the bar charts. Error bars indicate standard deviation, and student’s *t-test* was used to analyze statistical significance.

### Macrophage infection experiment

Metacyclic parasites were incubated with macrophages (J774A.1) for infection as described previously [25, 26]. Each experiment was carried out with three biological replicates and the data presented in the bar charts show the mean of three experiments, with error bars representing standard deviation. The two-tailed student *t*-test was applied to analyze statistical significance of the obtained data, and *p* values are mentioned in figure legends.

## Acknowledgements

We thank Ms. Pallavi Gulati for help in cloning the *set1* gene. We thank Dr. Vinay Nandicoori for extending the use of his laboratory facilities to us. DNA sequencing, confocal microscopy, and flow cytometry analyses were carried out with the help of instrumentation available at the Central Instrumentation Facility, University of Delhi South Campus.

## Data Availability Statement

All relevant data are within the manuscript and its Supporting Information files. Ld1S *set1* sequence has been deposited in GenBank. Accession no: OR479702.

## Supporting Information

**S1 Table. TUNEL Assay.** The data from two independent experiments have been tabulated. Percent values are average of two experiments.

**S1 Fig. Analyis of LdSET1 amino acid sequence in comparison with amino acid sequences of SET1 of other trypanosomatids.** Clustal Omega analysis viewed using Jalview multiple alignment editor. SET and post-SET domains are demarcated with black and red boxes respectively. Colors indicate the physico-chemical properties of the amino acids. Pink-hydrophobic/aliphatic; red-acidic; purple-glycine / proline; yellow-cysteine; orange/ochre-aromatic; dark blue-basic; green-hydrophilic.

**S2 Fig. A. Schematic representation of the domains** carried by LdSET1. **B. Western blot analysis of whole cell lysates isolated from Ld1S transfectant promastigotes,** probed with anti-FLAG antibody (Sigma Aldrich, 1:1000 dilution). Loading control: tubulin. **C. Immunofluorescence analysis of SET1-FLAG at different cell cycle stages** using anti-FLAG antibody (1:100 dilution). N: nucleus. K: kinetoplast. Kinetoplast morphology and segregation pattern was used as cell cycle stage marker. One roundish/short kinetoplast, one nucleus (1N1K): G1/ early S phase. One elongated kinetoplast, one nucleus (1N1K): late S/ early G2M phase. Two nuclei, one kinetoplast (2N1K): G2M phase. Two nuclei, two kinetoplasts (2N2K): post-mitosis. Magnification bar: 5 μm.

**S3 Fig. Analysis of expression of LdSET1 in *L. donovani* parasites. A.** Western blot analysis of whole cell lysates isolated from 4×10^7^ logarithmically growing and stationary phase promastigotes using anti-SET1 antibodies (already available in the lab; 1:2500 dilution). Loading control: tubulin. **B.** Western blot analysis of whole cell lysates isolated from 4×10^7^ procyclics and metacyclics using anti-SET1 antibodies. Loading control: tubulin. **C.** Western blot analysis of whole cell lysates isolated from synchronized *L. donovani* promastigotes (7×10^7^ cells per time-point). Upper panels: Flow cytometry profiles depicting the different cell cycle stages at which extracts were isolated. Lower left panels: western blots using anti-SET1 and anti-SET29 antibodies (already available in the lab; 1:2500 dilution). SET1 was maximally expressed in S phase while SET29 was equivalently expressed at all stages. Loading control: tubulin. Lower middle and right panels: quantification of expression of SET1 and SET29 in western blots of three experiments, by Image J analysis. Average values are plotted, error bars depict standard deviation, student’s *t*-test was applied for statistical significance. \**p* value < 0.05, \*\**p* value < 0.005, \*\*\**p* value < 0.0005, ns not significant.

**S4 Fig. Creation of *set1^-/+^***. Recombination at both ends was checked by PCRs across the deletion junctions. Positions of primers used are indicated with arrows. F1, F2, F3, F4: forward primers. R1, R2, R3: reverse primers. PCNA served as input template DNA control. Lanes 1: Ld1S genomic DNA template. Lanes 2: *set1^-/+^* genomic DNA template. M: DNA ladder.

**S5 Fig. Creation of *set1^-/-^***. Recombination at both ends was checked by PCRs/ inverse PCRs across the deletion junctions. For inverse PCR analyses, genomic DNA was digested with AvrII and SpeI enzymes (positions marked on the line diagram) and self-ligated. Positions of primers used are indicated with arrows. F3, F4, F5: forward primers. R1, R2, R3, R4: reverse primers. PCNA served as input template DNA control. *set1*: PCR for *set1* gene. Lanes 1: Ld1S genomic DNA template. Lanes 2: *set1^-/+^*genomic DNA template. M: DNA ladder.

**S6 Fig. A. Effect of higher concentrations of H_2_O_2_ on parasite survival.** *set1*-null cultures were initiated at 1×10^6^ cells / ml, and hydrogen peroxide (500 or 1000 μM) added on reaching a cell density of ∼ 7-9 x10^6^ cells / ml (Day 6). Cells were counted every 24 hours thereafter. **B. Effect of H_2_O_2_ on subcellular localization of SET1-FLAG.** *set1*^+/+^ cultures were initiated at 1×10^6^ cells / ml, and hydrogen peroxide (100μM) added on reaching a cell density of ∼ 7-9 x10^6^ cells / ml (Day 3). Subcellular localization was analyzed after 5h exposure to H_2_O_2._ Magnification bar: 5 μm.

**S7 Fig. Analysis of DNA damage in *set1^+/+^*and *set1^-/-^* cells.** Parasites were treated with 200 μM H_2_O_2_ for 5 hours and analyzed by TUNEL reaction. Each panel depicts merged image of fluorescein-UMP labelled DNA and DAPI fluorescence. Magnification bar: 10 μm.

